# A Large Yield Model for Crop Production and Design in Western Canada

**DOI:** 10.64898/2026.04.08.717277

**Authors:** Jordan Ubbens, Prachi Loliencar, Sateesh Kagale

## Abstract

With a changing climate, disease pressure, and other production threats, it is critical to ensure that crop producers are well-positioned to protect and optimize yields. In this work we present LYM-1, the first large-scale, multi-crop model for the prediction of yield performance in the Canadian prairies. This is enabled by a large dataset containing over 4.7 million yield observations across 10 different crop types, distributed over 23 growing years. Leveraging additional data sources for weather and soil properties allows the model to reason about the complex interactions between genetics, environment, and management which underlie yield. The trained model is not only effective at predicting the yield for held-out data, but also reveals scientifically and agronomically relevant effects such as the interaction between solar radiation and nitrogen uptake. We show that the architecture and training approach used in LYM-1 allows it to perform inference given incomplete queries where some model inputs are unspecified, while also establishing an advantage over popular linear and machine learning models in predictive performance.

## Introduction

Crop yields are complex, determined by a mixture of interacting factors, including solar radiation, precipitation, extreme weather events, genetic potential, fertilizer, irrigation, pest control, and inoculation, among many others (Yamoah et al., 1998; Lobell and Field, 2007; Assefa et al., 2018). Crop productivity also faces a period of high uncertainty due to the varied effects of climate change (Hultgren et al., 2025). With a complicated landscape of factors threatening crop yields, accurate and comprehensive models of yield are imperative for producers and an emerging tool for plant breeders.

The prairie provinces are a key region for food production in Canada. The province of Saskatchewan is one of the most important agricultural drivers, responsible for over half of the country’s oilseed production and up to 81% of its durum wheat production (St. Pierre and Mhlanga, 2022). The agricultural sector is also critically important to Saskatchewan’s provincial economy, accounting for over 7% of the province’s gross domestic product (Statistics Canada, 2025). For these reasons, crop productivity is vital to the prairie region’s economic success, the Canadian export market, and global food security.

Much recent and historical research has focused on the prediction of crop yields, and on estimating the relative effects of different environmental and management variables. One successful approach involves process-based physiological crop models which use forward simulation to model the biological processes which drive biomass accumulation and yield. Popular software packages have been developed for this type of functional crop modelling, including DSSAT (Jones et al., 2003), APSIM (Holzworth et al., 2014), and many others. These models explicitly represent underlying physiological processes including phenology, photosynthesis, water and nutrient uptake, and carbon allocation, allowing them to capture responses to abiotic stresses such as drought and heat, as well as management factors such as fertilizers. Comparisons among models indicate that explicit physiological representation generally improves predictive performance over empirical approaches, although misspecification may pose a problem when extrapolating beyond the calibration data (Zhao et al., 2014).

Other yield modelling approaches aim to learn patterns from a dataset of yield observations, unconstrained by models of biological or physiological processes. Linear regression and its penalized variants, such as ridge regression and LASSO, have been widely applied due to their high level of interpretability (Shastry et al., 2017). However, these classes of models are not expressive enough to approximate complex, nonlinear interactions between arbitrarily many factors affecting yield. In these scenarios, linearized relationships tend to under-fit by averaging over complex local interactions in the data, potentially missing agronomically meaningful effects (Ansarifar et al., 2021). Most contemporary models are based on machine learning techniques, leveraging the flexibility and representational capacity of these approaches to fit the complexities of the training data. Decision tree–based ensemble methods, including random forests and gradient boosted decision trees, have gained prominence due to their ability to model nonlinearity, along with their strong predictive performance on tabular data and a moderate level of interpretability (Van Klompenburg et al., 2020). Deep neural networks for yield prediction are also common in the recent literature, mirroring their popularity in other fields (Kick et al., 2023; Oikonomidis et al., 2023). Unlike linear methods and classical machine learning techniques, deep neural networks specialize in learning powerful representations from extremely large amounts of data. These strengths come at the cost of decreased interpretability as well as poor generalization performance when trained on small amounts of data, due to the tendency of overparameterized models to overfit.

Past attempts to build yield models based on deep neural networks have been limited by the size and scope of the data (Oikonomidis et al., 2022; Goel and Pandey, 2024). The raw quantity of the data is one of the most important factors when training high-capacity models, but there are also several quality characteristics which are equally important. In order to learn a comprehensive model of yield, the training data needs to incorporate a large number of growing years which represent a wide variety of climate regimes such as drought years. The data also needs to be geographically diverse in order to cover a wide range of soils and local microclimates. However, increased geographical and temporal scale can create other quality issues which cause deleterious biases in the data. For example, when one covariate is collected from multiple sources, different methodologies in measurement and differences in instruments may create artifacts in the data akin to batch effects which arise from these unobservable confounders. Ideally, the data should not only be large, but also free of unobserved sources of bias.

Here we describe LYM-1, a transformer-based deep neural network which is trained on a large multi-crop dataset containing over 4.7 million yield observations from the Canadian prairie region. In this work, we describe three novel contributions. First, we demonstrate that a transformer model trained with masked language modelling criteria enables the model to approximate the conditional distribution among tokens, allowing the model to make a prediction about any variable conditioned on an arbitrary subset of other variables. This means that the model is able to make predictions when prompted with partially complete queries, where many variables are left unspecified. Second, we show that such a model trained on a large corpus of yield observations outperforms linear and tree-based models on predicting the yield in unseen growing scenarios. Finally, we show that the same model can make plausible conditional predictions under partial queries, in general agreement with effects already established in the agronomic literature. The model is available for public use at https://bioinfo.nrc.ca/lym.

### Dataset Description

In this work, we use a crop insurance dataset collected by the Saskatchewan Crop Insurance Corporation (SCIC) to provide a fine-grained picture of the factors which influence crop productivity. We augment this data with weather data from Daymet (Thornton and Devarakonda, 2024), as well as soil data from the Canada Land Inventory (Agriculture and Agri-Food Canada, 1998). Table 1 summarizes the variables present in the data and their various sources, while Table 2 shows a summary of the dataset by crop type. Yield observations across time and space are shown in Supplementary Figures 4 and 5.

**Table 1.**
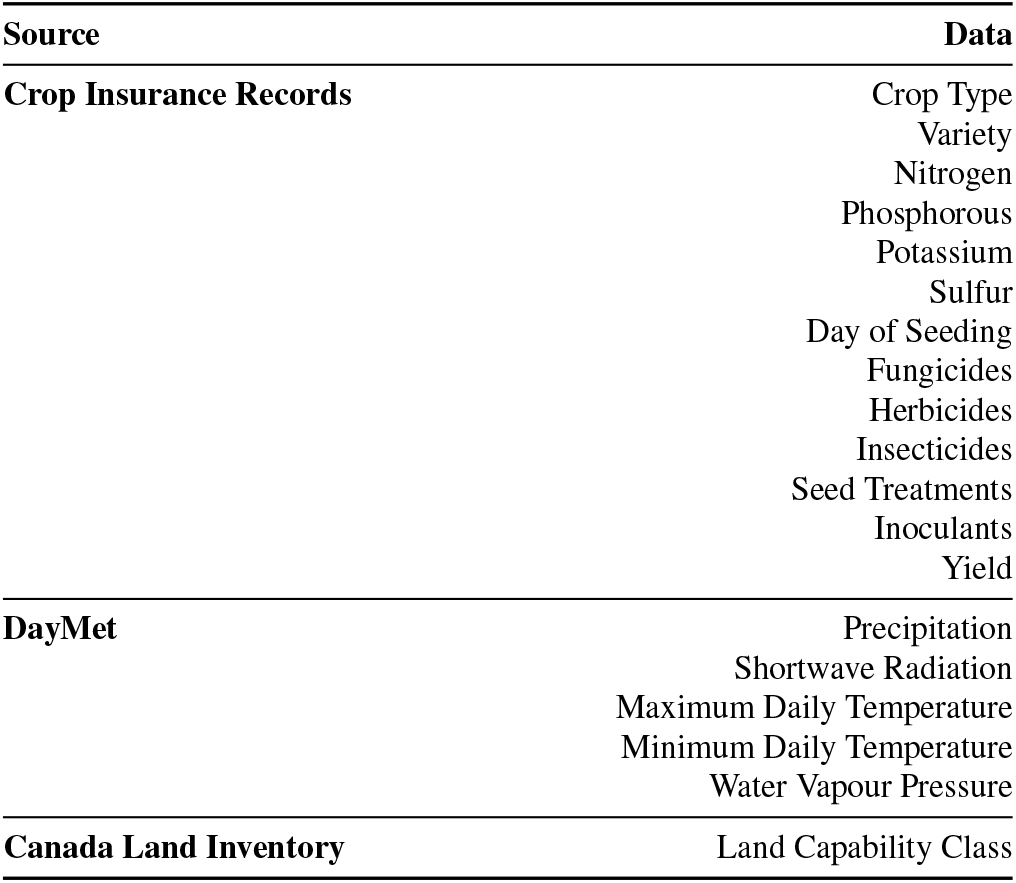
Summary of variables and sources of data used in the model.

**Table 2.** Summary of the dataset by crop type.

| Crop | Observ. | Years | Varieties |
| --- | --- | --- | --- |
| **Canola** | 1,421,014 | 1998 - 2021 | 713 |
| **Faba bean** | 3,729 | 1998 - 2021 | 15 |
| **Field Peas** | 431,265 | 1998 - 2021 | 197 |
| **Lentils (Red)** | 213,061 | 2001 - 2021 | 41 |
| **Lentils (Large Green)** | 146,897 | 2001 - 2021 | 21 |
| **Lentils (Other)** | 53,082 | 2001 - 2021 | 33 |
| **Wheat (Durum)** | 861,661 | 1998 - 2021 | 55 |
| **Wheat (Hard Red Spring)** | 1,466,820 | 1998 - 2021 | 168 |
| **Wheat (Hard White Spring)** | 4,448 | 2006 - 2021 | 8 |
| **Wheat (Winter)** | 32,653 | 1998 - 2021 | 30 |

The dataset includes a legal land description (LLD) associated with each yield observation, describing the geographical location of the field. A grid system is used to divide the province into land parcels called sections; each quarter of a section, approximately 160 acres, corresponds to a unique LLD. Shapefiles of the grid and QGIS were used to obtain the latitudes and longitudes of the the centroids of each LLD. These points were used as representatives for the LLDs.

Weather data for field locations was obtained from the DayMet weather service (Thornton and Devarakonda, 2024), which provides the daily weather measurements for the corresponding DayMet pixel (one square kilometer) of each LLD. In total, 5 weather features were collected: daily precipitation, shortwave radiation, maximum and minimum daily air temperature, and water vapour pressure. One feature, day length, is correlated with latitude and so it was excluded to avoid inadvertently exposing the model to any location information. Another feature, snow water equivalent, was not used as it is not specified during the growing season. See Thornton and Devarakonda (2024) for more details and methodology.

To assess the influence of the soil quality on yield, the Canada Land Inventory, published by the Agriculture and Agri-food Canada, was used. This inventory categorizes Canadian soil into eight classes based on agricultural ability: 1) no limitations for agriculture, 2) moderate limitations, 3) moderately severe limitations, 4) severe limitations, 5) forage crops (improvement practices feasible), 6) forage crops (improvement practices not feasible), O) organic soils, and n) unclassified. QGIS was used to sample these maps at the representative centroids of each LLD.

For the variety information, the date of registration was obtained from the Canadian Food Inspection Agency’s database of registered varieties. Varieties were anonymized using unique codes for the company and variety name.

For chemical inputs, the active ingredients were extracted from the product label found on the manufacturer’s website. For some legacy products, if the product label was not available online, the active ingredients were extracted from academic publications in the agronomy literature. This allows the inputs to be unified under the same identifier if they contain the same combination of active ingredients, even if sold under different product names. Each unique combination of active ingredients was encoded as a unique identifier. Chemicals were encoded as combinations instead of being broken out into the individual ingredients due to potential interaction effects between multiple ingredients when applied together in mixture. All 620 chemical products present in the raw data were mapped to an active ingredients grouping. 115 of these groupings harmonize more than one product, while a further 192 were singular 1:1 mappings of a single product to a unique grouping. Although a best effort was made to correspond products to active ingredients, this is a manual process and misclassifications are possible. A summary of the effect of each type of chemical input on yield is shown in Supplementary Figure 6. For each chemical, the effect size (Cohen’s *d*) between the entries where the chemical input was applied and where it was not was calculated. To ensure a sufficient sample size, only chemicals where there was a minimum of 90 entries where the input was applied are represented. For each chemical input, the yield data was orthogonalized by fitting an OLS regression model on the fertilizer data, precipitation, solar radiation, maximum and minimum temperatures, vapour pressure, as well as the remaining chemical data. This was done to control for the effect of weather as well as correlations between the application of chemical and fertilizer inputs.

The variables in the final dataset showed differing degrees of missingness. Weather and soil data were available for almost all datapoints, while variety and day of seeding were specified for 63% and 65% of the rows, respectively, and fertilizer information available for about 28%. A detailed breakdown of missingness is shown in Supplementary Figure 1.

#### Fertilizers support yield with diminishing returns

Producers balance the costs of agronomic inputs with the benefits realized in terms of yield and yield stability. Therefore, it is of interest to model the response of various crop types to fertilizers in order to optimize producers’ returns on investment. Figure 1 shows the response curves for nitrogen in canola and hard red spring wheat. The yield values were orthogonalized by fitting a linear regression using phosphorous, potassium, and sulfur, as well as precipitation, solar radiation, maximum and minimum temperatures, and vapour pressure, and obtaining the residuals from this model. This helps to control for weather differences as well as addressing collinearity between fertilizers, as when multiple inputs are applied, the amounts applied tend to be correlated. A polynomial of degree 3 was then fit on these residual values (blue line). The first derivative of this polynomial was calculated (dark grey line). Intuitively, when the derivative is above zero (shown by the light green shaded area), adding more of the compound increases yield, and when the derivative is below zero, adding more is deleterious to yield. The dark green vertical line indicates the maximum of this derivative, denoting the point of diminishing returns in the case of a parabolic response.

**Fig. 1.**
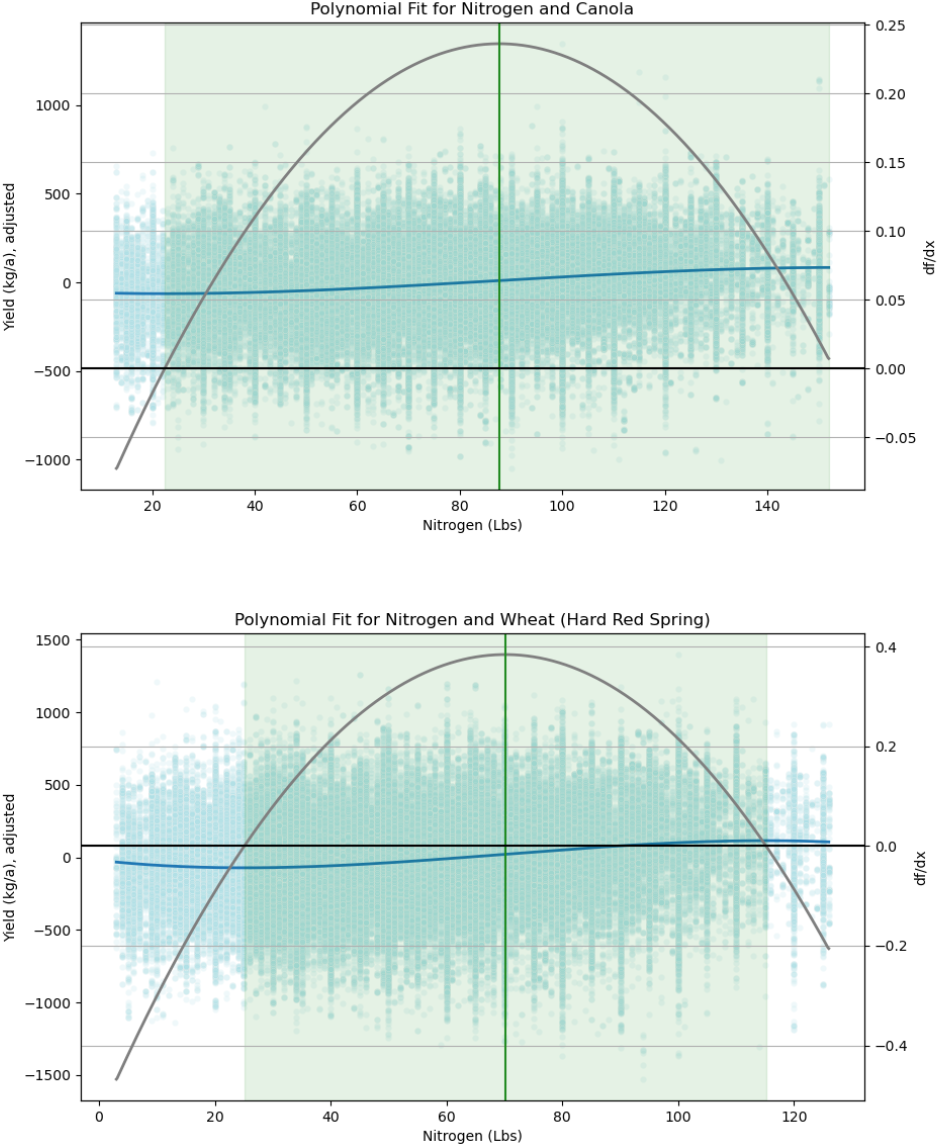
Example response curves for Nitrogen in canola (top) and hard red spring wheat (bottom). The range where applying more of the input results in increased yield is shown in green, and the point of diminishing returns is shown by the vertical line.

All fertilizers showed a parabolic type response where the effectiveness of each compound accelerated after a critical point, before dropping off after a attaining a certain concentration. Canola benefited from a significantly higher level of nitrogen, with a point of diminishing returns at around 85-90 pounds versus durum and hard red spring wheat at around 60-70 pounds. Both crops showed similar response to phosphorous, with diminishing returns reached at around 20-25 pounds and deleterious effects after about 30 pounds.

#### Better crop varieties have improved yields over time

Across crop plants, breeding has been fundamental in developing new varieties which are capable of delivering enhanced yield potential, as well as resistance to production threats such as drought and disease. To evaluate progress in variety development over time in Saskatchewan, yield performance was analyzed for registered varieties across wheat, lentils, canola, and field peas. After removing outliers and removing varieties with fewer than 90 samples to ensure good variance estimates, fertilizers, precipitation, and shortwave radiation were controlled for as well as minimum and maximum temperatures, aggregated weekly over the course of the growing season (weeks 14 to 38). The effect size with respect to yield between each variety and all others was then calculated using Cohen’s *d*. By plotting the effect size against the year of registration, it is shown that varieties have improved across all crop types in terms of their yield potential (Figure 2).

**Fig. 2.**
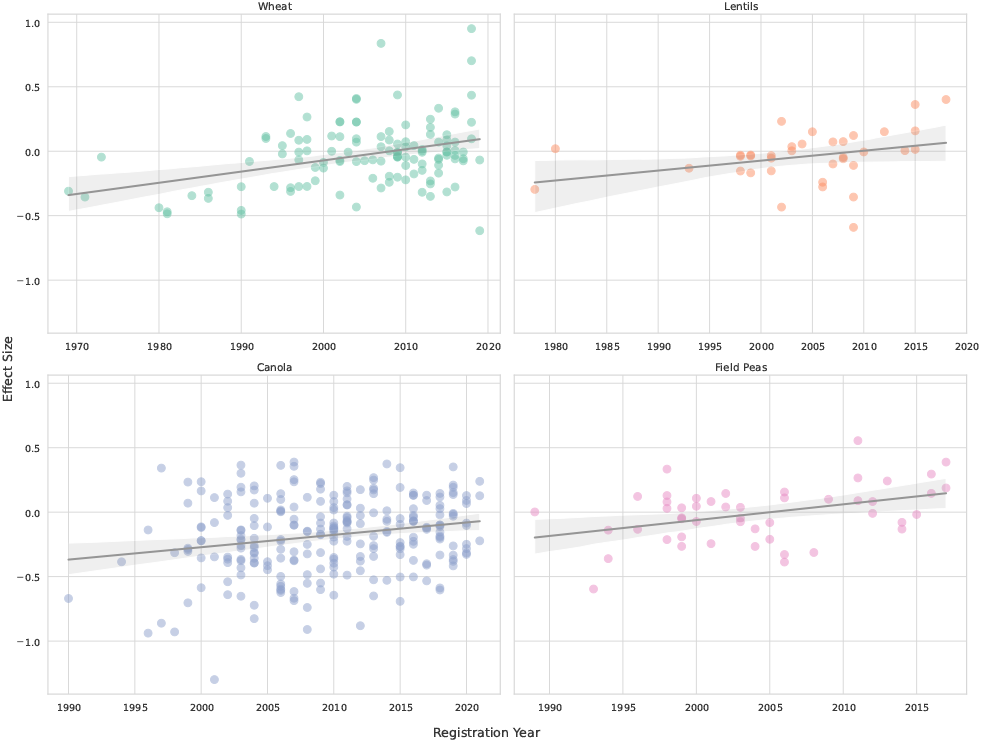
Effect size on yield for registered varieties of wheat, lentils, canola, and field peas versus the year of registration. Newer varieties have improved yields over time, controlling for weather and fertilizer inputs.

This analysis captures only the raw difference in yields, but there are other breeding objectives which are not represented here, such as yield stability, water use efficiency, or resistance to new disease threats. There may also be some varieties which, for example, were primarily recommended for substandard soil types, affecting the overall yield outcome. Therefore, this should be taken as only one component of genetic gain over time, among many. There may also be other management trends which vary over time, affecting the yield trend independently of weather and fertilizer use.

### Creating a Large Yield Model

We describe a Large Yield Model called LYM-1 which enables detailed inference in arbitrary growing scenarios. This model aims to approximate the full conditional distribution, using the high representational capacity of deep neural networks. The remainder of this section describes the methodology for training the model.

#### Preprocessing

A preprocessing pipeline was developed to prepare the data prior to pretraining. The daily weather data was aggregated by taking the mean over months or weeks during the growing season (from April to September). This allowed for the development of two different model variants with different levels of granularity for weather. Next, entries with zero kilograms of production were removed. These zero entries could be present for various reasons, but in fact may represent non-zero yields which were classified as total losses due to their monetary value being below the cost of harvesting. For numeric variables, outlier removal was performed by identifying datapoints which exceeded two times the inter-quartile range, and the data were rescaled to between -0.5 and 0.5. Standardization was not used due to the fact that some variables were zero-inflated and many were multi-modal or otherwise non-normal. For the non-weather continuous variables, outlier removal and rescaling were both performed independently on a per-crop basis, due to the fact that crop types tend to be distinct from one another in various ways. For example, a particular seeding date may be extremely early for pea, but common for winter wheat. The weather variables were rescaled, but no outlier removal was performed.

#### Pretraining

The model architecture and pretraining process are based on masked language modeling (MLM) (Devlin et al., 2019; Liu et al., 2019), using an encoder-only transformer (Vaswani et al., 2017). In order to adapt MLM to heterogeneous token types (such as weather, chemical inputs, etc.), each token is passed through a type-specific embedding module and then conditioned with a token type embedding. The embeddings *t*_*i*_ are input to an encoder-only transformer model, which provides output embeddings *e*_*i*_. These output embeddings are then decoded using a token type-specific decoder. Linear projections were used for all decoders and for the encoders of continuous values. An overview of the full model is shown in Figure 3.

**Fig. 3.**
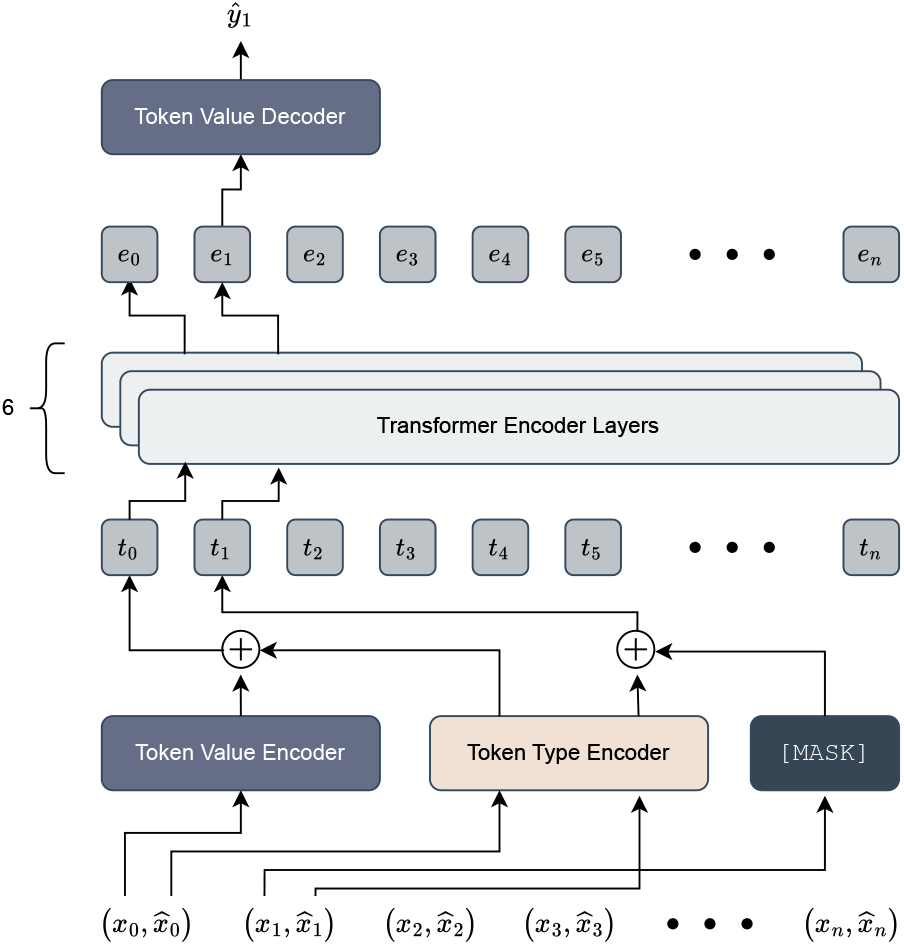
General architecture for the Large Yield Model. The input pairs 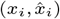 represent the value and the type of input, respectively. In this example, the second input token 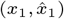 is being predicted.

During pretraining, the model is provided with token value and token type pairs 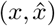. The value *x* is randomly replaced with the mask token with probability *p* = 0.5. A learning rate schedule with a linear warm-up and a cosine decay is used, with a maximum learning rate of 1 *×* 10^−4^ and a batch size of 128. AdamW is used as the optimizer with *β*_1_, *β*_2_ = 0.9, 0.999. A standard encoder-only transformer is used with an internal dimensionality of 1024, 16 attention heads, Pre-LN (Xiong et al., 2020), and a total of 6 encoder layers, for a total of approximately 75 million non-embedding parameters. Hyperparameters were tuned by hand on the one-epoch experiments, using training loss as a proxy for validation loss due to the single epoch. Performance on the validation set was never monitored during training and early stopping was not used.

During training, the L2 loss is used for the masked continuous tokens and cross-entropy loss is used for the masked categorical tokens. The loss is weighted using an uncertainty-based multi-task loss weighting (Kendall et al., 2018). The loss is therefore given by

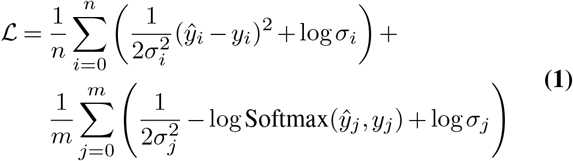

where (*ý*_*i*_, *y*_*i*_), *i* ∈ 0 … *n* represents the targets and predictions for the *n* masked continuous tokens, (*ý*_*j*_, *y*_*j*_), *j* ∈ 0 … *m* represents the targets and predictions for the *m* masked categorical tokens, and *σ* are the learned variance parameters for each token type.

#### Finetuning

In order to specialize the pretrained model for yield prediction, finetuning was explored using two different techniques. Each proceeded from a pretrained model which was pretrained for four epochs, with the yield token removed entirely. Optimization during finetuning used the same learning rate schedule as pretraining with a maximum learning rate of 2 *×* 10^−4^. The first finetuning technique used a mean pooling of the outputs from the second to last transformer encoder layer, resulting in 1024-dimensional features. This representation was fed into an MLP regression head with a hidden dimension of 512, directly predicting yield. The entire transformer was frozen and weight updates were applied to the regression head only. As an alternative, CLS finetuning was also explored. During finetuning, a CLS token was appended to the input sequence. A linear projection from the output token was used to predict the yield label. The first two transformer encoder layers were frozen, and the remaining four, as well as the regression head, were updated during training. Freezing the initial transformer layers and updating the subsequent ones is common practice in CLS finetuning for natural language tasks (Ingle et al., 2022). It theoretically allows the model to maintain the same initial representations learned during pretraining, which are expected to be robust general representations, while using the remainder of the layers to model the downstream task. It also accelerates pretraining since fewer parameters are updated.

## Results

Here, we test the ability of LYM-1 to produce accurate yield predictions in unseen scenarios, test its ability to reproduce effects which are known to impact yield, and explore some hypotheticals by crafting novel queries.

### Validation Performance

To evaluate the ability of LYM-1 to generalize to unseen data, a set of yield observations was held out as a validation set. This validation set was used to assess the predictive performance of both pretrained models as well as finetuned models. For the pretrained models, the yield token was included during pretraining and replaced with a mask token during inference. In addition to the 75M parameter model, a larger 126M parameter model was tested for pretraining by extending the number of layers from six to ten. A summary of predictive performance can be seen in Table 3. Detailed predictions and residuals for a pretrained model and a finetuned model are shown in Supplementary Figures 2 and 3.

**Table 3.** Validation performance for pretrained and finetuned models. Weather is specified monthly. MAE indicates mean absolute error, RMSE root mean squared error, R^2^ the coefficient of determination, and *ρ* Spearman’s rank correlation. PT: pretrained, MP: finetuned via mean pooling, CLS: finetuned via CLS token.

| Parameters | Type | Epochs | MAE | RMSE | $R^2$ | $\rho$ |
| --- | --- | --- | --- | --- | --- | --- |
| **Pretrained Models** |  |  |  |  |  |  |
| 75M | PT | 1 | 0.092 | 0.118 | 0.50 | 0.69 |
| 126M | PT | 1 | 0.089 | 0.115 | 0.53 | 0.72 |
| 75M | PT | 2 | 0.087 | 0.113 | 0.54 | 0.71 |
| 126M | PT | 2 | 0.088 | 0.113 | 0.55 | 0.73 |
| **Finetuned Models** |  |  |  |  |  |  |
| 75M | MP | 1 | 0.084 | 0.111 | 0.56 | 0.73 |
| 75M | MP | 2 | 0.084 | 0.110 | 0.57 | 0.74 |
| 75M | CLS | 1 | 0.081 | 0.106 | 0.60 | 0.76 |
| 75M | CLS | 2 | 0.076 | 0.101 | 0.64 | 0.76 |

The model demonstrates an ability to predict yield in unseen data, whether it is finetuned on yield after pretraining or simply pretrained with the yield variable included. The highest predictive accuracy for pretrained models was seen when using the larger 126M-parameter version and training for two epochs, with an R^2^ of 0.55 between predicted and actual yield. When finetuning the smaller 75M-parameter models, CLS finetuning proved more effective than finetuning using the mean pooling representation, with an R^2^ of 0.64 when training for two epochs compared to 0.57 for the latter.

In Figure 4, LYM-1 is compared to several prediction baselines. Here, mean (crop) indicates predicting the mean of the respective crop (null performance because yields are roughly zero-centered on a per-crop basis) and mean (crop-year) indicates predicting the mean of the respective crop during the respective year. Year is not provided to the models so mean (crop-year) is not directly comparable. For ridge regression and random forest we use the implementation from scikit-learn 1.9.0 (Pedregosa et al., 2011), and for XGBoost we use the official implementation (3.4.1) (Chen and Guestrin, 2016). For ridge regression and random forest, missing data is not supported and so missing fields were imputed. Continuous variables were imputed based on the crop-specific median value and a special missing category was added for categorical variables. XGBoost supports missing data natively, and so these values were left as null. Hyperparameter tuning was performed for each baseline model using 5-fold cross-validation on the training set. The full hyperparameter grids are shown in Supplementary Table 1. This analysis shows a clear advantage for the proposed method, even compared to methods such as random forests and XGBoost which have a history of strong generalization with tabular data. The fine-tuned version of LYM-1 outperforms the next best performing method (XGBoost) by over 39%. It should be noted that the model has not been extensively optimized for validation performance, so there is likely more headroom for widening the performance advantage over the baseline models. For example, it is sensible that using a finetuned model with a larger number of parameters can increase validation performance (extrapolating from Table 3).

**Fig. 4.**
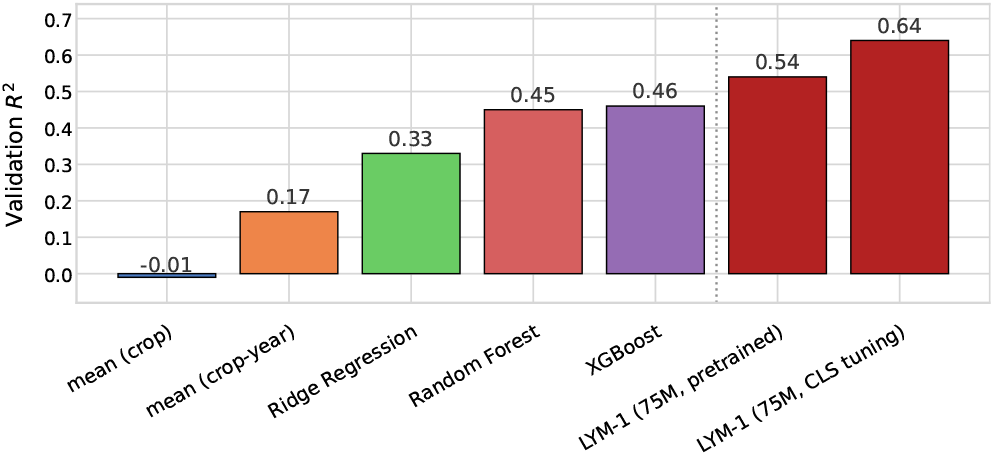
Validation performance of the 75M parameter pretrained and finetuned models relative to baselines.

### Ablations

The results presented in Table 3 represent the predictive performance of the model on unseen datapoints drawn uniformly from the dataset. However, it is also necessary to investigate the generalization performance of the model on out-of-distribution data. For this reason, we performed two ablation tests using validation sets which were out-of-year and out-of-region. The results of these experiments are shown in Table 4.

**Table 4.** Generalization experiments for different held-out validation sets. The 75M model with CLS finetuning for one epoch is used. Weather is specified monthly. MAE indicates mean absolute error, RMSE root mean squared error, R^2^ the coefficient of determination, and *ρ* Spearman’s rank correlation.

| Validation Set | MAE | RMSE | R<sup>2</sup> | $\rho$ |
| --- | --- | --- | --- | --- |
| Uniform (Table 3) | 0.081 | 0.106 | 0.60 | 0.76 |
| Held out 2021 (drought) | 0.089 | 0.115 | 0.43 | 0.70 |
| Held out southern 25% | 0.099 | 0.127 | 0.40 | 0.61 |

For the out-of-year experiment, we left out 2021 and trained on the remaining years (1998-2020), resulting in a validation set of 182,271 samples. This represents a difficult split, as 2021 was an extreme drought year and so the model needed to predict yields far below the mean of its training set. For the out-of-region experiment, we held out all datapoints in the lower 25% of the geographical region, based on the recorded latitude, resulting in a validation set of approximately 1.35 million samples. This represents a similarly difficult split of the data, as the province shows a high degree of difference in geography and climate along this axis, from boreal forest in the north to sand desert in the south.

Although the performance of the model decreases on these deliberately difficult train-test splits, it retains significant predictive performance with an R^2^ of 0.40 to 0.43. Some of the difference between these results and the uniformly sampled validation set (R^2^=0.60) may be due to differences in the size of the training sets, but we believe that these splits demonstrate the worst-case scenario for generalization with the data available.

In addition, the effect of specifying weather aggregated weekly or monthly during the growing season was tested, with negligible differences in prediction performance (Table 5). This may indicate that intra-month differences have only a small predictive effect compared to smoothing over the full monthly period.

**Table 5.**
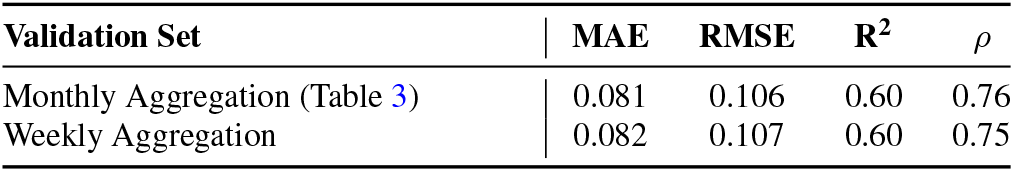
Comparison of predictive performance with weather aggregated monthly versus weekly. The 75M model with CLS finetuning for one epoch is used. MAE indicates mean absolute error, RMSE root mean squared error, R^2^ the coefficient of determination, and *ρ* Spearman’s rank correlation.

### Reproducing Known Effects

In the quantitative results, LYM-1 proves capable of predicting yield performance in unseen data. However, the utility of the model depends on its ability to provide accurate conditionals in the presence of incomplete data. In the previous evaluations, the model was provided with all of the available data, including measurements for most of the available token types, such as weather, variety, fertilizers, and so on. However, when predicting the effects of individual differences, many of the possible token types may be left undefined by the user. This means that these inputs will be out of distribution for the finetuned models, which expect a full set of observations. Simply imputing values for these missing token types at inference time is impossible, as the full joint distribution between token types is unknown. However, using the pretrained models in this way is well-motivated as they are trained to predict the full conditional distribution with multiple missing token types caused by masking. This can be done at inference by simply adding any token type not specified by the user (except for chemical inputs) as a masked token, in order to keep the inputs in-distribution. Although finetuned models attain the highest performance in validation, using a pretrained model is the best strategy for handling incomplete inputs during end-user testing.

Interrogating the model in this way reveals interesting inferences. Here, we first conduct a series of smoke tests to check whether what the model has learned about the architecture of yield coincides with prior knowledge. For example, Figure 5 shows the inferred differences among land capability classes for hard red spring wheat. This shows the expected ordering, where higher-quality soils are predicted to provide higher yields across seeding dates, although the model predicts a marginal narrowing of these gaps between soils at later seeding dates.

**Fig. 5.**
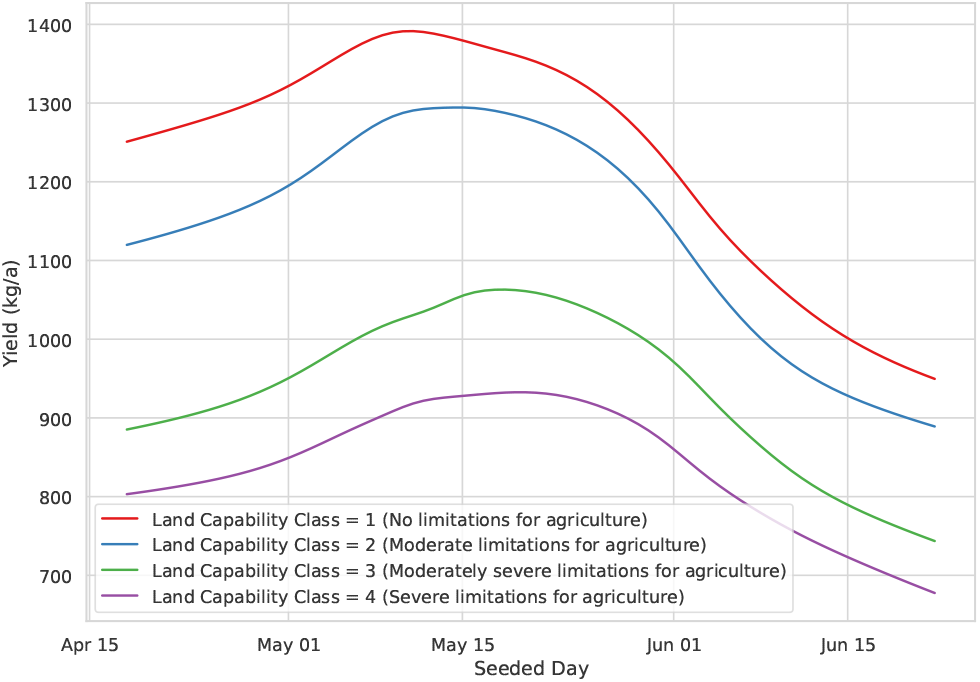
Comparison of the inferred effect of different land capability classes on yield for hard red spring wheat. Other specified variables: crop type: hard red spring wheat, soil capability class: 1.

It has been shown that July is one of the most important times during the growing cycle for canola, where excess heat has an adverse effect on yield (Kerr et al., 2019). Figure 6 shows that the model infers on the order of about 50 kilograms per acre loss due to an increase in the daily maximum temperature of about 1°C during July, regardless of seeding date. For the following analyses, land capability class was set to 1 (no limitations for agriculture) to control for differences in soil quality. Importantly, this analysis allows the model to infer other variables which are left undefined, such as precipitation, which may be correlated with temperature.

**Fig. 6.**
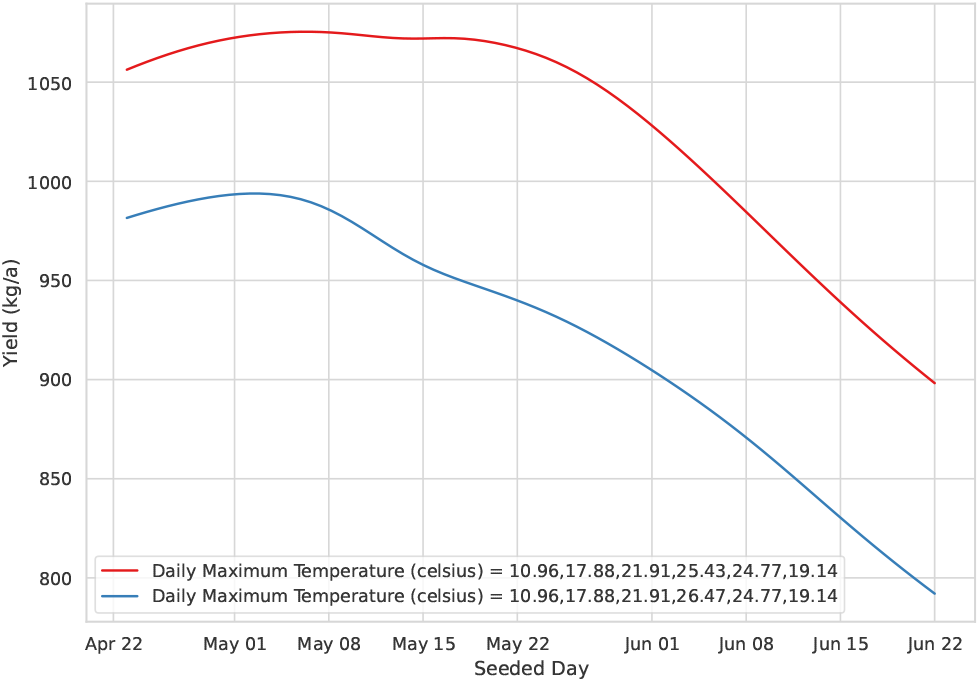
Inferred effect on canola yields of increasing daily maximum temperature by approximately 1°C during July (blue series), compared to the median temperature for all months (red series). Other specified variables: crop type: canola, soil capability class: 1.

The model is also capable of inferring interaction effects, by varying two axes simultaneously. Figure 7 shows the interaction between added nitrogen and shortwave radiation. The difference in inferred nitrogen use efficiency between moderate and extreme levels of solar radiation are in concordance with observations that light levels affect both the rate of nitrogen uptake as well as allocation of nitrogen resources within the plant (Sakuraba and Yanagisawa, 2018; Liang et al., 2022).

**Fig. 7.**
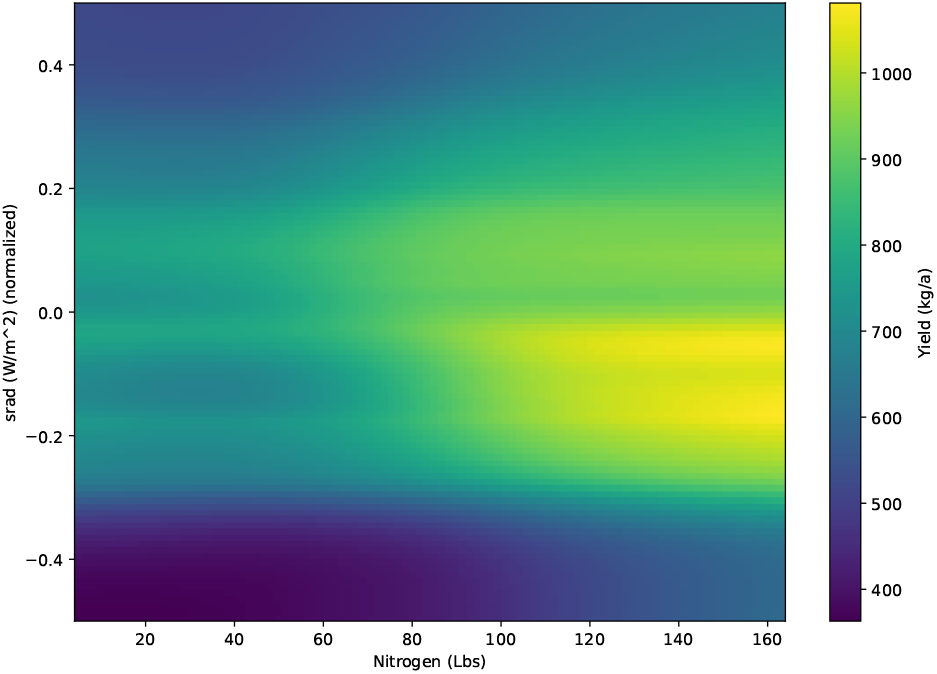
Inferred interaction effect between Nitrogen (pounds added) and shortwave radiation (W/m^2^) in canola. For shortwave radiation, the distribution for each time-point was scaled to between -0.5 and 0.5. Other specified variables: crop type: canola, soil capability class: 1.

### Testing Hypotheticals

Beyond recapitulating known effects, the model can also be used to make *in-silico* comparisons which would require extensive field trials to validate.

For example, comparing three of the most common fungicide mixes used in field peas shows differentiation in Figure 8. In this scenario, the model predicts that only the mixture of azoxystrobin and propiconazole results in a net increase in yield, with the other two having negligible or even detrimental effects compared to the baseline of no chemical inputs (red).

**Fig. 8.**
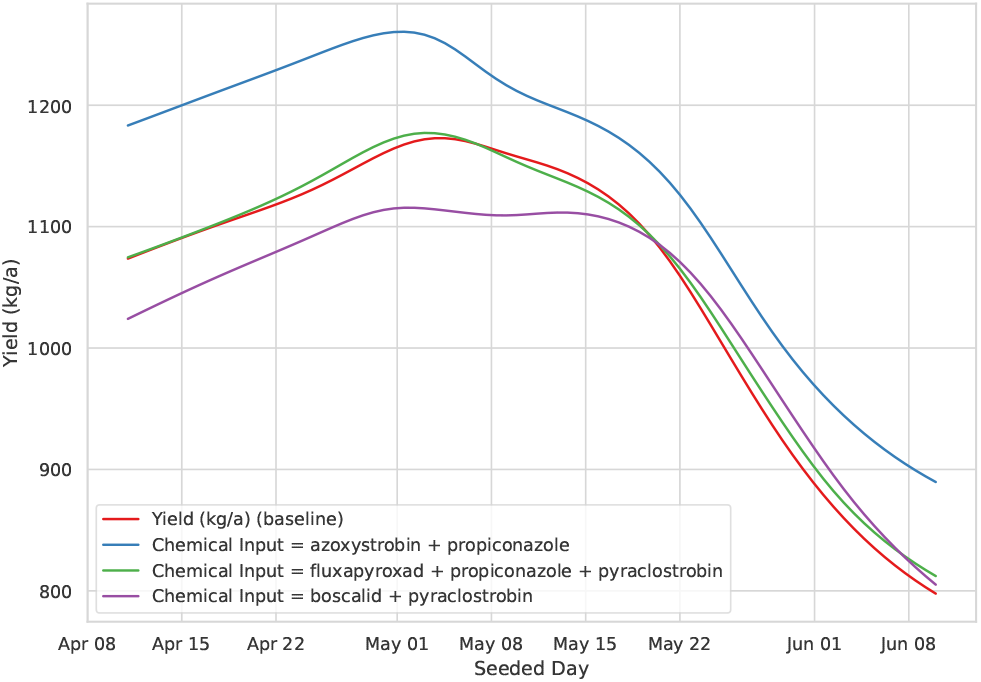
Comparison of inferred effects of three fungicide mixes in field pea. Other specified variables: crop type: field peas, soil capability class: 1.

The left panel of Figure 9 shows the yield performance of a variety of durum wheat registered in 2013 against a different line registered in 2019. All weather variables are unspecified. In the right panel, the average daily precipitation from the 2015 drought year is specified, as retrieved from the Saskatchewan Research Council Saskatoon climate reference station report ^1^ by dividing the monthly precipitation figures by the number of days in each month. This analysis shows how a durum wheat variety which was released in 2019 would have hypothetically performed during the drought of 2015. The 2019 line outperforms the 2013 line in general, but also shows a differential response with respect to the seeding date under drought conditions. The older line is roughly on par with the more modern line when seeded during May, but this parity slips in the drought condition and the newer variety is uniformly superior.

**Fig. 9.**
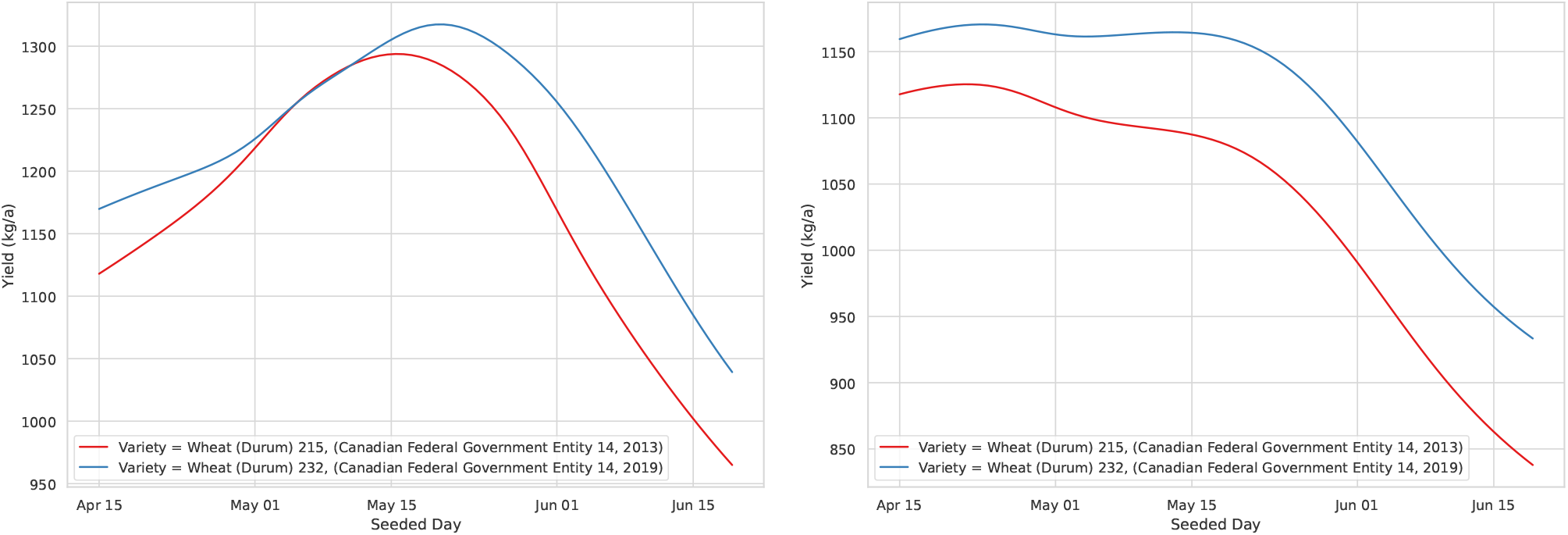
Inferred yield performance for two varieties of durum wheat under unspecified weather conditions (left), and under the precipitation conditions seen during the drought year of 2015 (right). Other specified variables: crop type: durum wheat, soil capability class: 1.

## Discussion

### Scaling

Even with 4.7M yield observations, LYM-1 likely remains data-limited. This is in contrast to large language models, which benefit from such a high volume of unstructured pretraining data that they are most often limited by available compute, not data. However, with a limited number of ablation experiments, we observe that LYM-1 seems to scale in compute in terms of both parameters and training time, even though the relatively small amount of data necessitates re-using data over multiple epochs (Table 3). The validation performance shown here can likely be increased by scaling up in compute, even with relatively limited data, although generalization performance is likely best improved through expanded training data.

### Limitations

Although LYM-1 proves to be a capable model for yield prediction, there are drawbacks to the approach and questions to be answered. One significant drawback is a lack of interpretability. Although some interpretability techniques do exist for transformer models (such as the attention maps shown in Supplementary Figure 7), interpretability in general remains a challenge for deep neural networks and other heavily nonlinear models. This is especially difficult in compute-heavy models where it is not tractable to re-train many versions of the model with different covariates. The lack of interpretability puts the approach in contrast to other methods in yield prediction which are based on generalized linear models or physiological crop models, which can use their advanced interpretability to verify the feasibility of the model. The approach of relying on a high-capacity nonlinear model to learn the dynamics of the data trades away this interpretability for representational capacity.

The strength of using the masked training objective is that the trained model can be given only a subset of variables during inference, while the rest are considered to be masked. This means that neither the model not the user needs to know the full joint distribution of the variables in order to perform inference. However, just as interpreting the outputs of the model is difficult, one must also be aware that the outputs reflect observational correlations which may not necessarily be causal in nature. For example, if the query specifies high nitrogen input and leaves temperature unspecified, if high nitrogen input is correlated with low temperatures, the model will provide yield predictions as if the temperature were low. This can be circumvented by also specifying temperature in the query, but it is important that users of such models have an awareness that incomplete queries do not necessarily mean that the unspecified variables have no effect on the predictions.

## Conclusion

Crop yields are complex, determined by a large array of interacting factors. Here we presented LYM-1, a large yield model designed to learn the architecture of yield from a large and diverse dataset of western Canadian field crops, assembled from multiple data sources. LYM-1 was shown to be capable of predicting yield performance in held out data across 10 different crop types. Qualitative testing revealed that it also reproduced expected agronomic effects, such as the deleterious effect of excess heat in July on canola and the influence of solar radiation on nitrogen uptake. Access to the full model is provided online.

The approach taken by LYM-1 demonstrates that, by increasing data size well into the millions of datapoints, state-of-the-art techniques in modelling become possible in the yield prediction space. Future models may simply expand the breadth of the training data beyond the single geographical region used in this work, into different types of climates, as well as the different crop types which are associated with those regions.

## ACKNOWLEDGEMENTS

We gratefully acknowledge the Saskatchewan Crop Insurance Corporation (SCIC) for providing the dataset used in this analysis. Loliencar was supported by the National Research Council Postdoctoral Fellowship program.

## Footnotes

1 https://www.src.sk.ca/sites/default/files/resources/2015%252520crs%252520stoon%252520annual%252520summary.pdf

## Notes

### Competing Interest Statement

The authors have declared no competing interest.

### Summary of Updates

Additional experiments for comparison to baseline models; ablations for out-of-distribution generalization performance; monthly versus weekly weather aggregation; revised supplementary materials.

