## Supplemental Materials for "A Large Yield Model for Crop Production and Design in Western Canada"

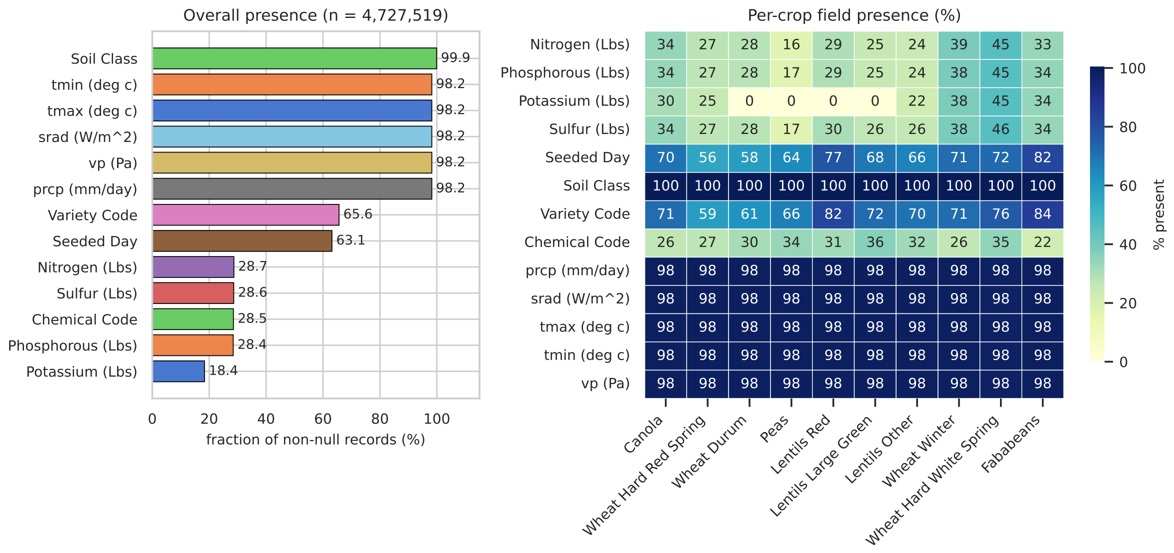


Supplementary Figure 1: Summary of missing data values (left) and a detailed per-crop view (right).

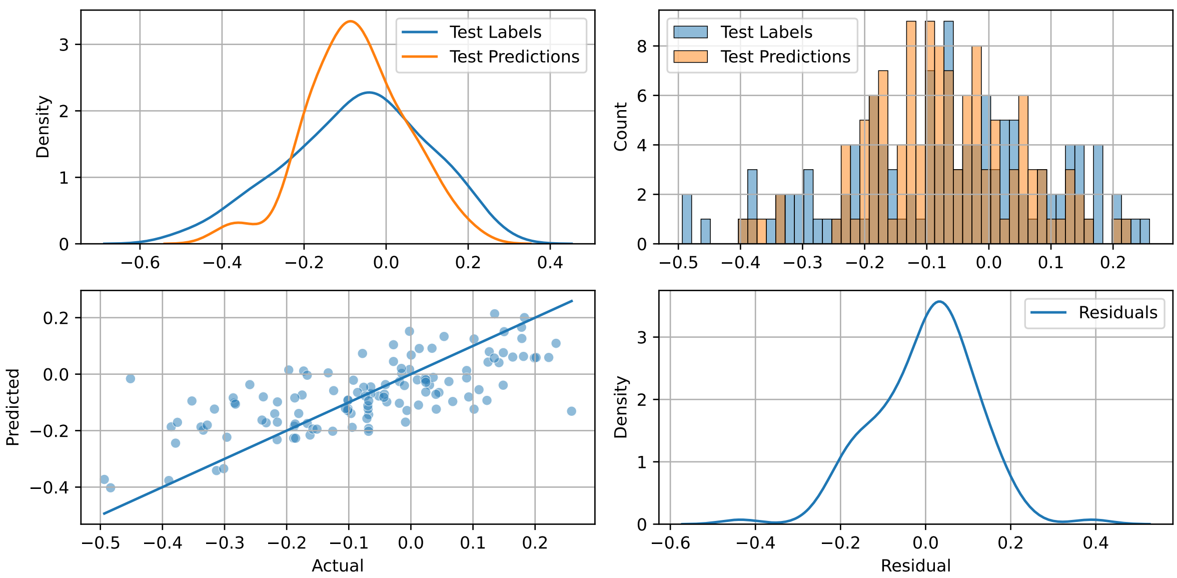


Supplementary Figure 2: Validation performance for LYM-1 (75M) pretrained model

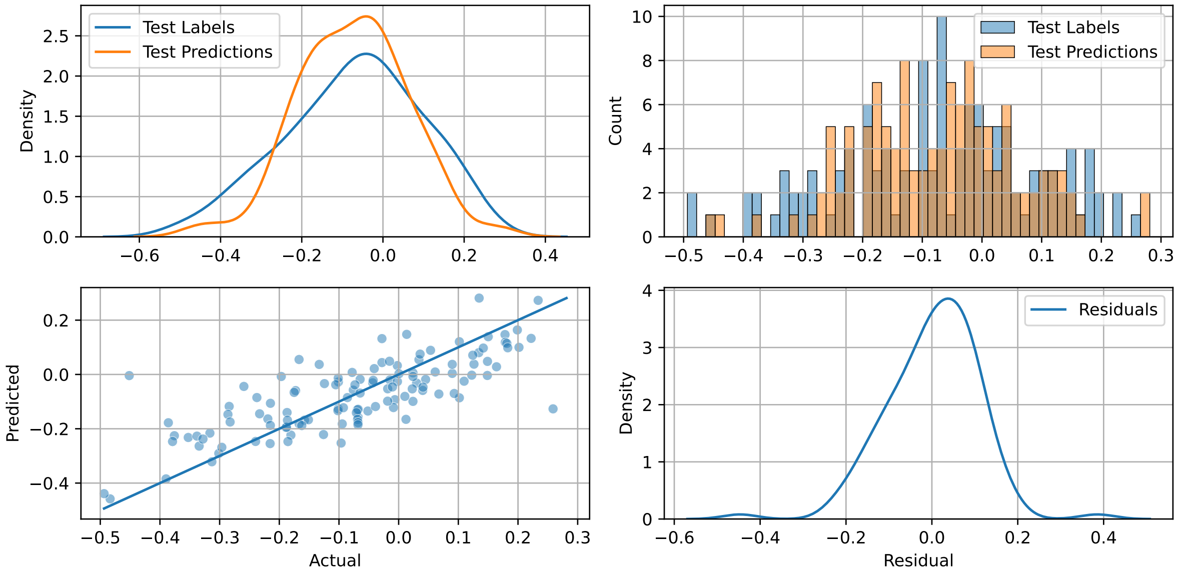


Supplementary Figure 3: Validation performance for LYM-1 (75M) CLS finetuned model

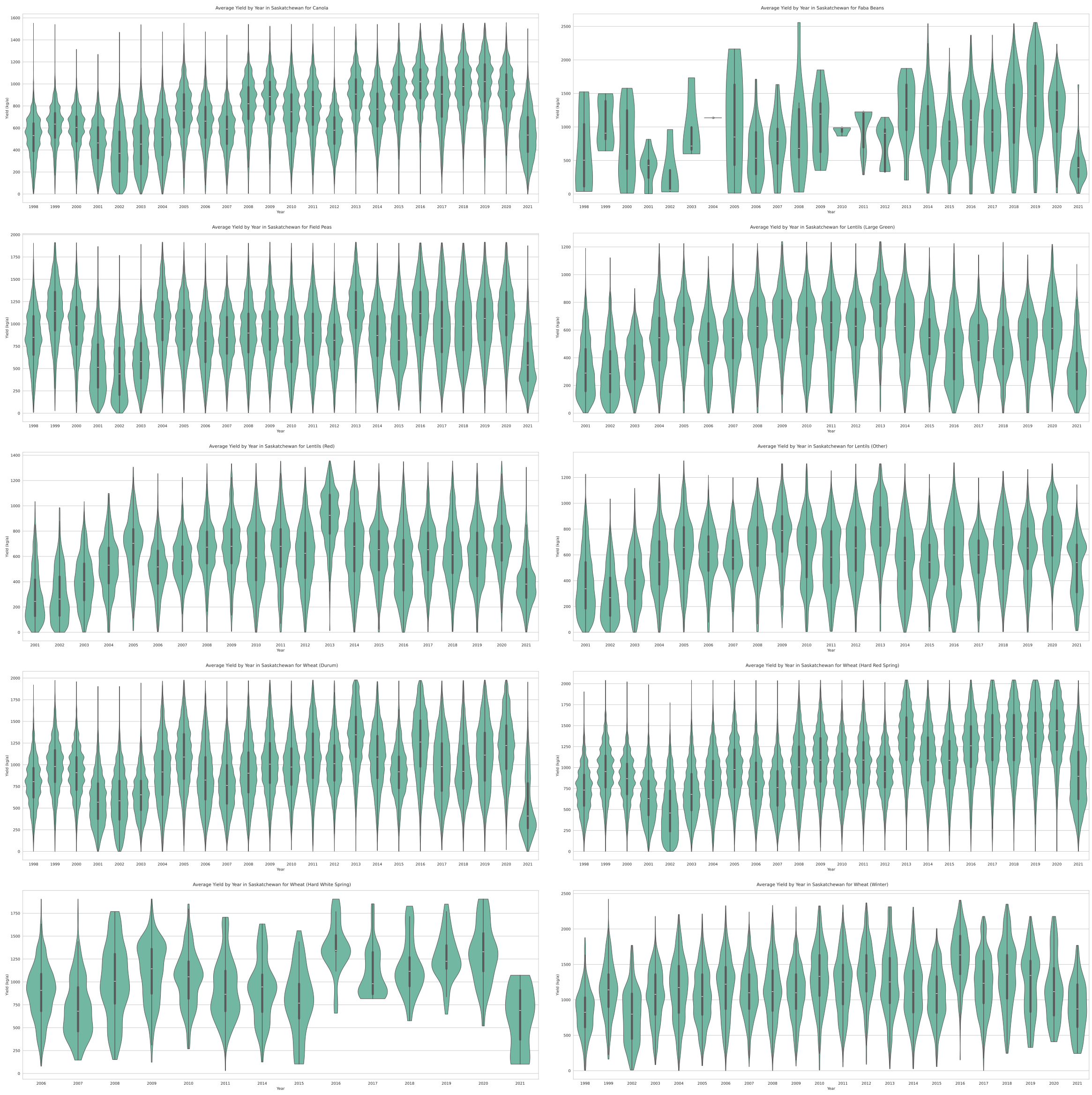


Supplementary Figure 4: Observed yields by year.

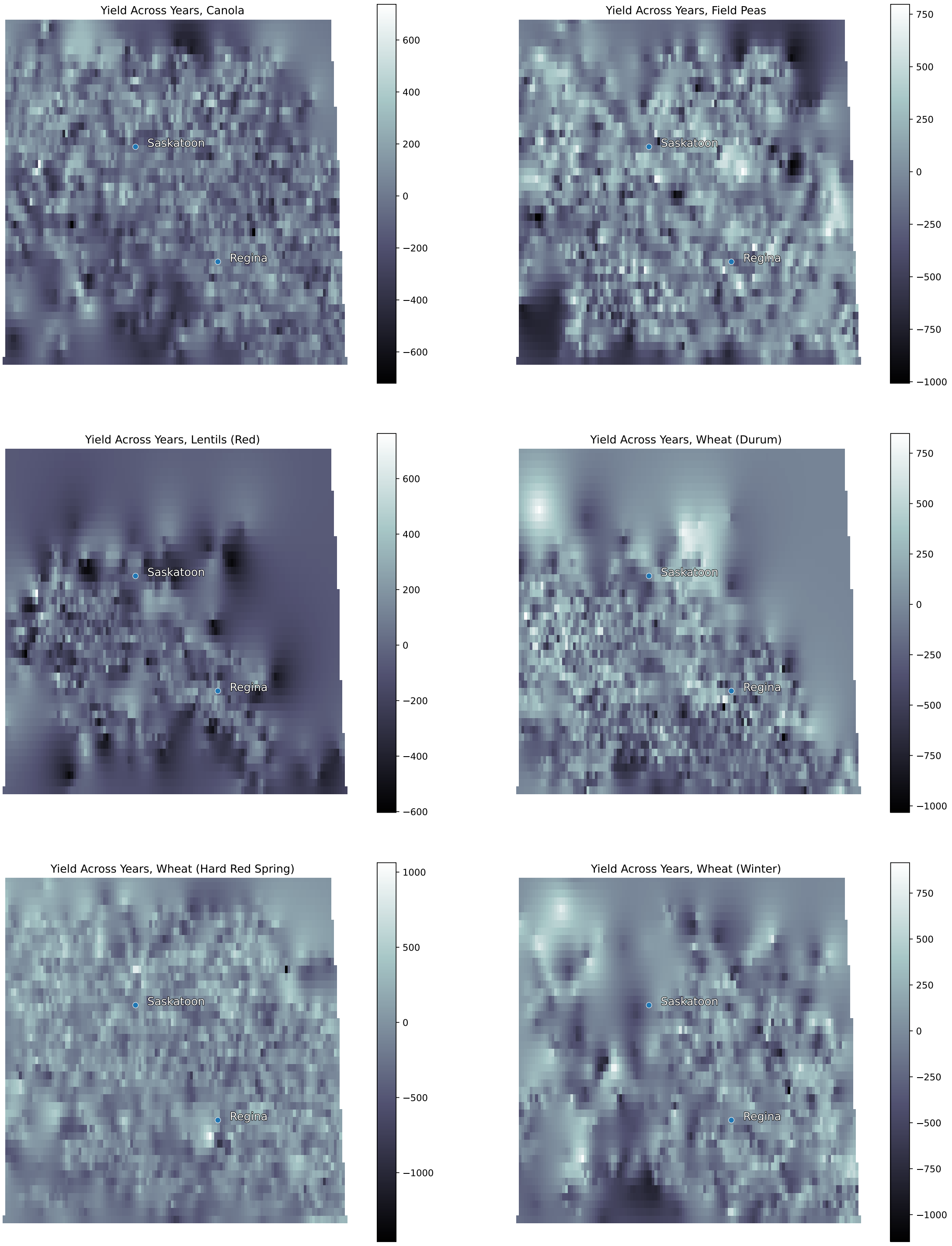


Supplementary Figure 5: Interpolated yield data for various crop types across the arable region of southern Saskatchewan. Chemical and fertilizer inputs were controlled for. The data was downsampled to 3000 observations using a spatially-aware downsampling method which thinned samples in areas of high point density. Kriging was used to interpolate the data spatially.

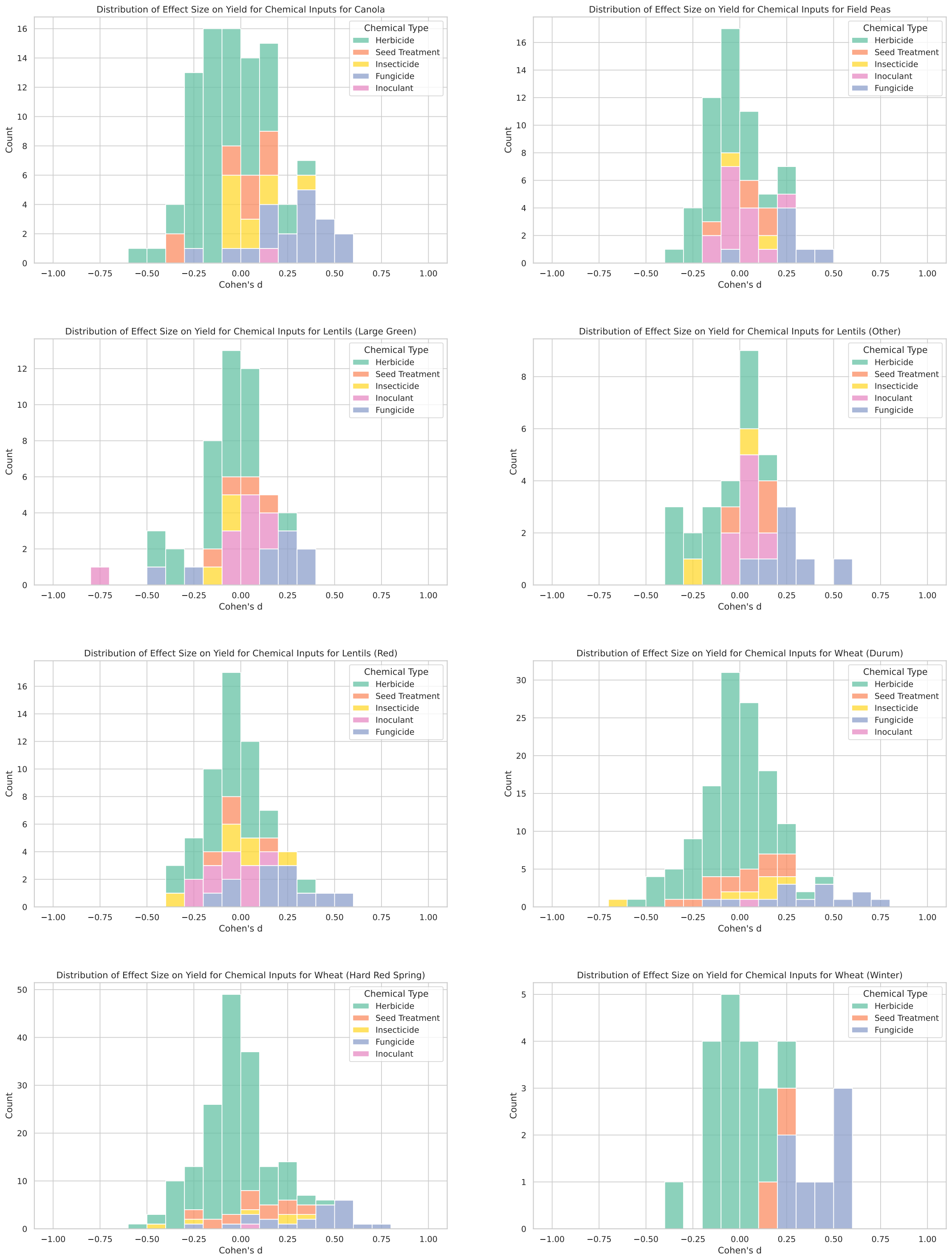


Supplementary Figure 6: The effect on yield of different categories of chemical inputs across crop types.

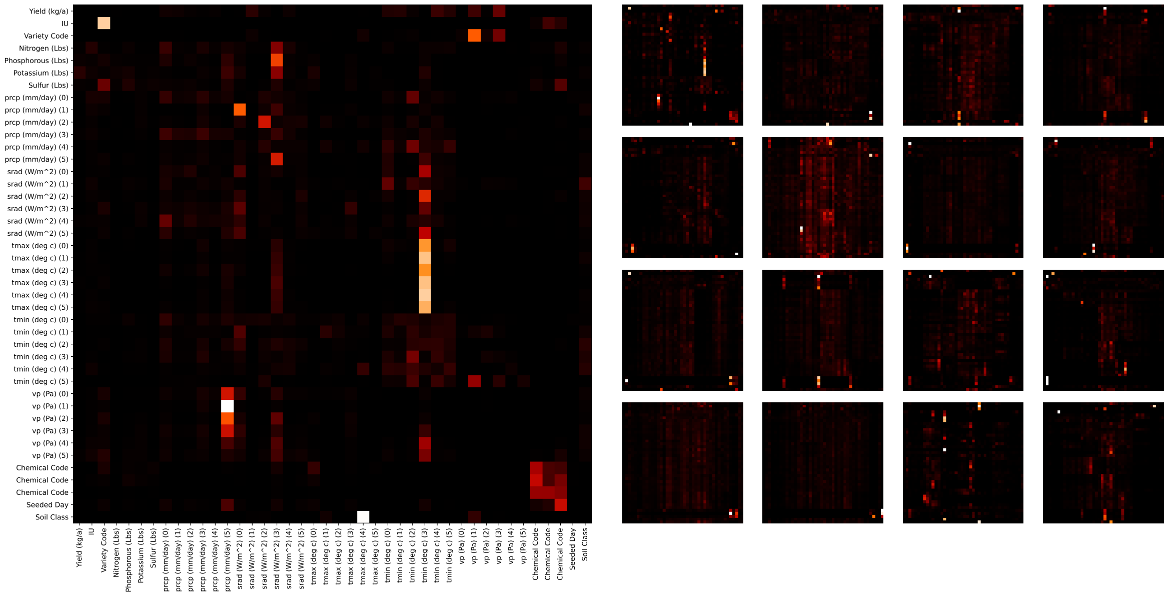


Supplementary Figure 7: Visualizing the attention maps for the 16 attention heads in the first layer (right). The first attention head is shown in more detail on the left. The input is a single sample from the validation set and the yield token is masked.

| **Method** | **Hyperparameter** | **Search values** |
| --- | --- | --- |
| Ridge Regression | $\alpha$ | $\{10^{-2}, \ldots, 10^{4}\}$ (logarithmic) |
| Random Forest | `n_estimators` | {50, 100, 150} |
| | `max_depth` | $\{3, 6, \infty\}$ |
| | `min_samples_leaf` | {1, 2, 4} |
| XGBoost | `n_estimators` | {50, 100, 150} |
| | `max_depth` | $\{3, 6, \infty\}$ |
| | `learning_rate` | {0.1, 0.2, 0.3} |

Supplementary Table 1: Hyperparameter grids used for baseline methods.
